# Towards a mental programming neural circuit: Insights from working memory sequence manipulation

**DOI:** 10.1101/2025.07.24.666222

**Authors:** Junfeng Zuo, Cheng Xue, Si Wu, Wen-Hao Zhang

## Abstract

Cognitive reasoning, also known as mental programming, is fundamental to our intelligence. A central function in mental programming is working memory (WM), involving both temporarily maintaining the information and manipulating it based on rules. While the neural mechanisms of WM maintenance have been extensively studied, those governing WM manipulation remain largely unknown. To bridge this gap, the present study focuses on an elementary operation in WM manipulation: re-ordering items in WM sequences. We propose a functional, biologically plausible neural circuit model that consists of two interconnected modules: a memory module composed of continuous attractor-based memory slots that store item features, and a control module sending gain-modulating commands to orchestrate specific operations in the memory module. The model successfully implements two-item swapping in a WM sequence, generating neuronal responses similar to recent primate experiments of WM sequence manipulation. By incorporating principles from the algebraic permutation group, we generalize the circuit model to accommodate more complex sequence manipulations. This math foundation reveals how arbitrary permutations can be decomposed into sequences of elementary swapping operations, which can be generated by a hierarchical tree-structured control circuit module. And the mutual inhibition within the control tree ensures that only one program is being executed at the same time. Our study establishes overarching connections among mental programming neural circuit models, neuroscience experiments, and abstract algebraic structure. These insights enhance our understanding of the neural underpinnings of cognitive reasoning and inspire the design of artificial cognitive systems.

## 1 Introduction

Cognitive reasoning refers to a wide spectrum of mental processes such as logic reasoning, problem solving, decision-making, language, etc. It is a fundamental cognitive function and lays the foundation of intelligence [1]. Back to the era of Alan Turing, it has been hypothesized that the cognitive reasoning shares similar fundamental principles with computer programming [2], analogous to a kind of mental programming. A central component of cognitive reasoning is working memory (WM) [3, 4], including *maintaining* information (memory) temporarily and *manipulating* this information (mental operation) based on some *rules*. Studying how neural circuits in the brain implement WM maintenance and manipulation is fundamental in neuroscience, which can help us understand the neural basis of mental programming, and may inspire the next-generation AI systems. Meanwhile, although ML research has developed neural networks for complex programming and language processing (e.g., [5–8]), most of these networks lack neurophysiological support or far from biological plausibility.

A large body of experimental and modeling studies focused on the neural basis of WM maintenance [10–12], suggesting that WM items are maintained with a slot-like representation where each item is stored in a specific memory slot with architecture as continuous attractor networks (CANs) [13–17]. However, the circuit architecture and algorithm of WM manipulation remain largely unknown. An elementary operation of WM manipulation is to re-order WM sequences (Fig. 1A, [18–20, 9]), e.g., swapping two items, which is also a primitive operation in computer programming. Hence, studying re-ordering WM sequences can provide insights into the neural mechanism of WM manipulations.

**Figure 1.**
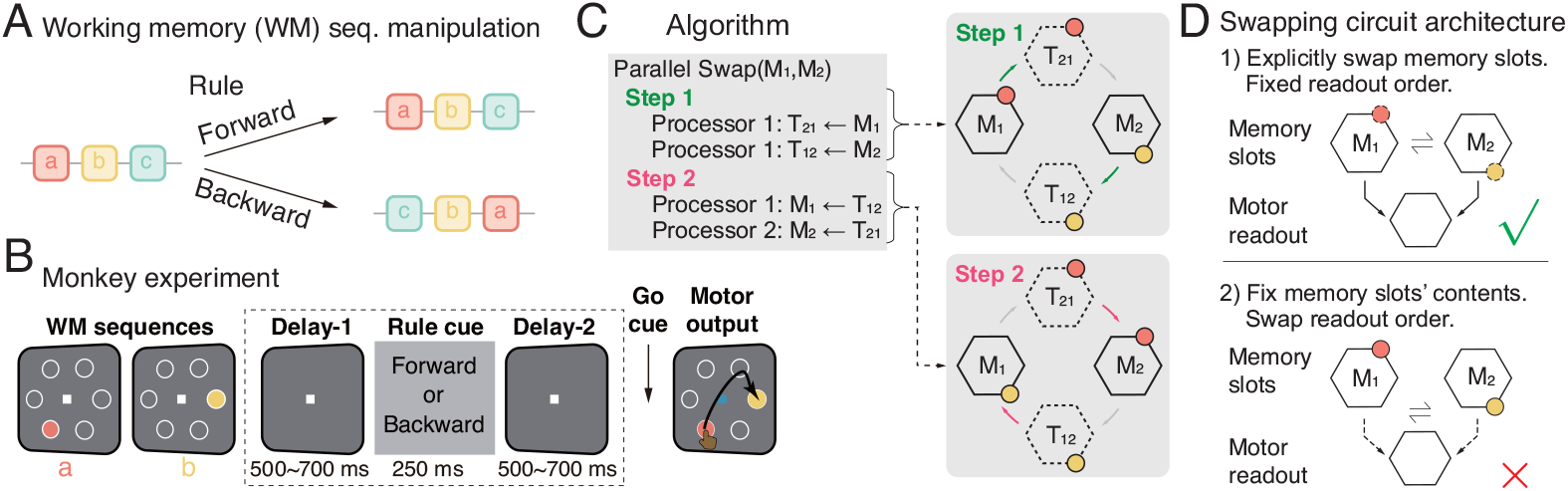
Working memory (WM) sequence manipulation is an elementary operation in mental programming. (A) The task of repeating or reversing a WM sequence. (B) Monkey experiments on WM sequence manipulation. The figure is adapted and modified from [9]. (C) The neural circuits’ parallel swapping algorithm found from the experiment in (B). (D) Two possible circuit architectures for the WM sequence manipulation. The experiment in (B) suggests the top one.

A recent neuroscience experiment on monkeys provided direct evidence for WM sequence reordering in frontal cortex neural circuits (Fig. 1B) [9]. It suggests that the frontal cortex employs an **architecture** explicitly swapping the WM items stored in two memory slots, and the motor output reads out these items from the slots in a fixed order (Fig. 1C-D). This rejects another possible architecture of implicit WM item swapping, where the motor readout is able to readout contents from memory slots with flexible orders (Fig. 1D, bottom). In addition, the experiments identified that the frontal cortex runs a **parallel swapping algorithm** with two steps (Fig. 1C): The first step is to load the contents from two memory slots to corresponding temporary slots, and then temporary slots are cross loaded back to memory slots, forming a circle structure among these four slots (Fig. 1C).

Inspired by this experiment, we propose a circuit model to maintain and manipulate WM sequences that consists of two interconnected modules: one is the memory module composed of continuous attractor network (CAN)-based (temporary) memory slots storing WM item features, and another is the control module issuing *gain-modulation* commands to orchestrate specific operations (programs) in the memory module. The control module consists of several sub-modules, each representing a program (rule) and commanding the memory module in a specific way. The circuit successfully swaps the two items in the WM sequence by utilizing the *autonomous* circuit dynamics, and generates neuronal responses comparable to experimental observations. Our model relies on three main circuit mechanisms: 1) It demonstrates that simple gain modulations, which are widely observed in the cortex, are able to coordinate complex operations in the memory module. 2) The inhibitory feedback from the memory to the control module confirms the execution of an operation, which is crucial for multi-step programs. 3) Mutual inhibition exists between memory slots and temporary slots, and across different control sub-modules to ensure that only one operation is executed at a time.

We further generalize the circuit for manipulations of longer WM sequences guided by the algebraic permutation group [21]. The re-ordering corresponds to a symmetric group, whose math representation suggests that a complex permutation can be decomposed into a series of elementary swappings. This directly specifies a tree-structured control circuit module for generating a sequence of swapping commands to realize a complex permutation. The construction of the hierarchical tree-structured control circuit is self-similar with the interaction between the swapping control module and the memory module, ensuring the scalability of the circuit architecture. As a proof of concept, we show a circuit model for 3-item permutations (*S*_3_ group) with a tree-structured control module.

### Significance

We propose one of the first biologically plausible neural circuit models for mental programming of WM sequence permutations, with solid experimental support and rigorous guidance from the algebraic permutation group. It implies that the algebraic group can be used to determine the neural circuit architecture, and sheds insight into neural networks for more complicated programming.

## 2 Towards a neurally plausible circuit model for mental programming

The present study takes a construction approach to build a biologically plausible circuit model for mental programming of WM sequence manipulations and motor readout, based upon well established canonical neural circuit motifs and neural operations. We will demonstrate that the known canonical motifs and operations can be organically combined to implement mental programming.

### 2.1 Circuit architecture overview and “design philosophy”

Since working memories involve the information maintenance and manipulation, the proposed WM circuit has two modules (Fig. 2A): One is the memory module storing the information, and another is the control module manipulating the WM content. In addition, the circuit has a third motor module that reads out the content in the WM circuit with a fixed order to generate (motor) outputs.

**Figure 2.**
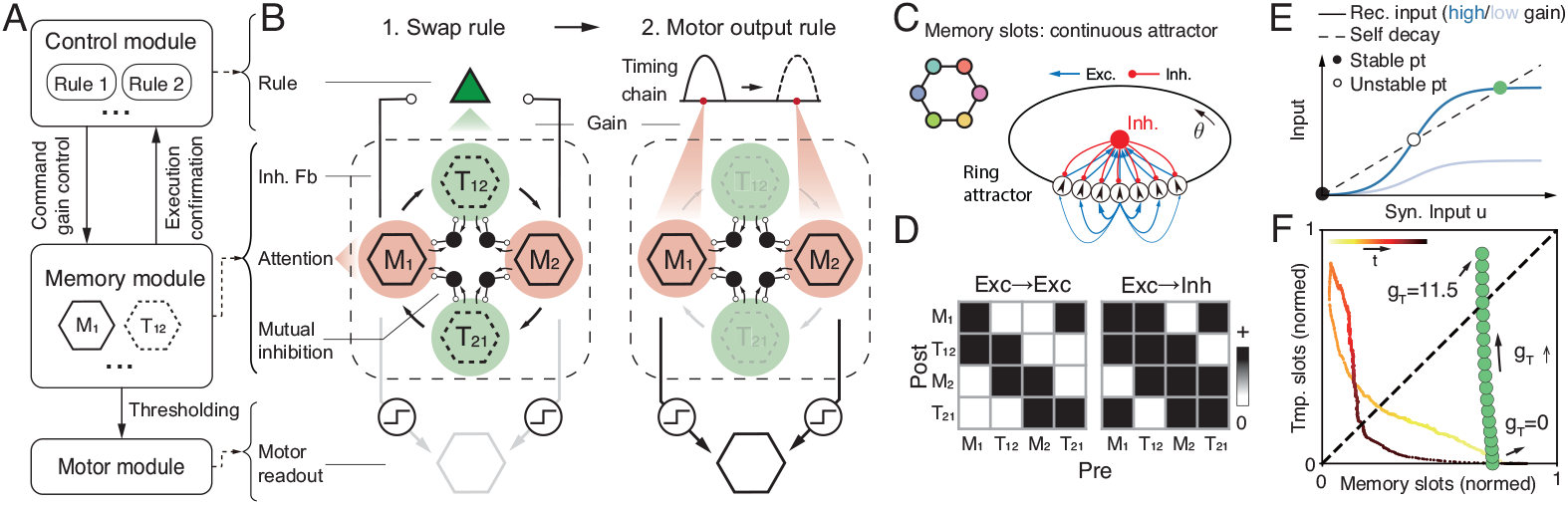
(A-B) The modular structure of the recurrent circuit for swapping and outputting two WM items. (C) Each (temporary) memory slot (*M*_1_, *M*_2_, *T*_12_, *T*_21_) is modeled as a continuous attractor network with excitatory and inhibitory neurons. (D) The connections between (temporary) memory slots. (E) The top-down gain control from the control module could switch ON (dark blue) and OFF (light blue) of memory slots. (F) The joint activities (total responses) of memory and temporary slots. Colored line: transient response during swapping; green dots: gain-dependent fixed points.

#### Interactions between WM modules

The memory and control modules are *recurrently* connected with each other, rather than just uni-directional connections from control to memory module (Fig. 2A). To increase the scalability of the model, the interaction between two modules is as simple as possible, and then we embed the computational complexity into the *autonomous* recurrent dynamics within each module. Specifically, we consider top-down *gain modulations* from the control to the memory module, which is a simple operation widely observed in the cortex without needing fine spatial structures at the single neuron level [22–24]. Meanwhile, the inhibitory feedback from the memory to the control module is crucial for the control circuit to register the execution of an operation.

#### Sub-modules of the control module

Depending on the operations (rules), the control module can be divided into several sub-modules with each providing different gain modulations to the memory module to run specific operations. For example, the WM swapping circuit has two rules (Fig. 2A): one is swapping WM items, and another is reading out WM sequences into the motor region.

### 2.2 Memory slots of stimulus features: continuous attractor networks

Mimicking WM experiments, each WM item is a continuous feature (e.g., direction in Fig. 1B), which needs to be stored in a structured memory slot. In the present study, the structured memory slots are modeled as CANs, which is a canonical circuit model in neuroscience researches on representing continuous features [13, 12, 25–28], and each of them can be regarded as a cortical hypercolumn. A CAN consists of excitatory (E) neurons and a pool of inhibitory (I) neurons (see detailed description in Sec. S1.1). The E neurons utilize their structured recurrent weights to give rise to structured population responses to store a WM item, while I neurons stabilize the network dynamics.

#### E neurons’ dynamics

E neurons are selective for a 1D angular feature *z* ∈ (− *π, π*]. Denote *θ*_*j*_ as the preferred stimulus feature of the *j*-th E neuron, and the preferred feature of all *N*_*E*_ E neurons, 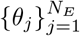, uniformly cover the whole space *z* (Fig. 2C). Mathematically, in the continuum limit of an infinite number of neurons (*θ*_*j*_ → *θ*), the dynamics of E neurons can be written as [29, 17],

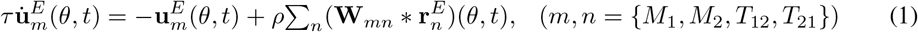

where 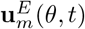 and 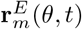 represent, respectively, the synaptic inputs and firing rates of neurons preferring *z* = *θ. m* is the index of memory slots that is referred to as one of the slots in Fig. 2B. *τ* is the time constant, and *ρ* = *N*_*E*_*/*2*π* is the neuronal density covering the stimulus feature space.

#### Recurrent connection kernel

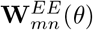 is the recurrent connection kernel from E neurons in memory slot *n* to the slot *m*, which are modeled as Gaussian functions in the model (Fig. 2C),

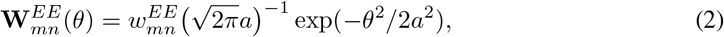

where 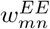 (scalar) is the peak recurrent weight and to be adjusted for realizing mental programming. *a* the connection width across the stimulus feature space. The symbol * denotes the convolution, i.e., **W**(*θ*) * **r**(*θ*) = **W**(*θ − θ*^*′*^)**r**(*θ*^*′*^)*dθ*^*′*^. The memory slots form a circle structure in connection topology (*M*_1_ → *T*_12_ → *M*_2_ → *T*_21_ → *M*_1_), with the connection weight shown in Fig. 2B and D.

#### Divisive normalization and gain modulation of E neurons

In a memory slot, the E neurons are subject to the global inhibition from a pool of I neurons, which is modeled as divisive normalization (activation function), a canonical operation observed in the cortex [30, 31].

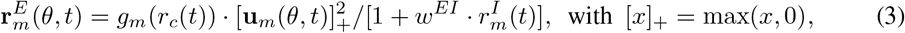

where 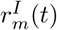 (scalar) is the instantaneous mean firing rate of I neuron pool (Eq. 4). *w*^*EI*^ (a positive scalar) characterizes the effective inhibition strength from I neurons to E neurons. Importantly, the E neurons are also subject to a gain *g*_*m*_(*r*_*c*_(*t*)) (positive scalar) modulated by the control circuit response *r*_*c*_ (Eq. 6). The gain can switch the circuits between *Up* (high response) and *Down* (low response) states via the cusp bifurcation (Fig. 2E) [32]. Therefore, the control circuit *r*_*c*_(*t*) (Eq. 6) will rely on gain modulation to command memory slots to run specific operations (see the Sec. 2.3-2.4). Moreover, the gain modulation is excitatory (*g*_*m*_ > 0) and *homogeneous* across all E neurons within a memory slot, which simplifies the interactions from control to memory module.

#### Shared inhibition across memory slots

Each memory slot has an I neuron pool without feature selectivity, so our model only preserves I neurons’ mean firing rates 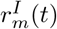 (scalar). The 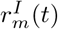 in a memory slot are driven by E neurons in the same slot and neighbor slots (Fig. 2B and D), which introduces competitions between memory slots that is crucial for autonomous swapping,

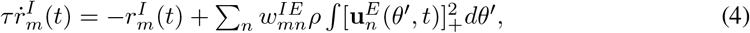

where 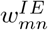 (scalar) is the weight from E neurons in slot *n* to the I neuron pool in slot *m*. All (temporary) memory slots in the circuits are *symmetric* with the same intrinsic parameters and connectivity. Their difference is the asymmetric gain modulation *g*_*m*_ (Eq. 3) from control circuits. The symmetric memory slots simplifies circuit architecture, improving its robustness and scalability.

### 2.3 The swapping control sub-module: gain modulation on (temporary) memory slots

A control circuit module receives the external rule signal of swapping (backward) and provides gain modulations to E neurons in the temporary slots, i.e., changing the *g*_*m*_ in Eq. (3), and meanwhile receives the feedback from memory slots. The mutual interactions between the control circuits and the memory module is crucial for utilizing the recurrent circuit’s **autonomous dynamics** to perform swapping while rule cues are transiently presented, reducing the complexity of control signals.

The swapping control circuit is also a recurrent circuit motif that consists of E and I neurons. Unlike the memory slots with structured recurrent weights between E neurons, the control circuit does not have item feature selectivity, and thus we only consider the mean firing rate of E neurons (scalar) in a control sub-module. To simplify the model further, we don’t explicitly model I neurons’ dynamics in the control circuit but absorb their effect into E neurons’ activation function (divisive normalization). Denote *u*_*c*_ and *r*_*c*_ as synaptic input and firing rate of E neurons in the control circuit respectively,

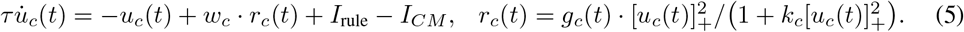

*I*_rule_ is the external rule signal, and *I*_*CM*_ is the inhibitory feedback from memory slots. The control circuit modulates the gain of E neurons in the temporary slots,

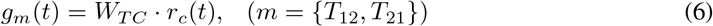

where *W*_*TC*_ denotes the weight from swapping control module to temporary WM slots.

### 2.4 The mechanisms and procedures of automatic WM item swapping

We explain the circuit mechanism for swapping content over memory slots, especially how the information only flows for half circle and then stops through autonomous dynamics coordinated by the gain modulation from the control circuit (Fig. 2B). The key mechanism is the gain modulation could flexibly switch ON and OFF of the information flow over memory slots, without changing the synaptic weights. This is different from the mainstream artificial neural networks that gate the synaptic weight matrix to route information flow [5, 6], where the biological plausibility of synaptic gating remains an issue while the gain modulation is ubiquitous across the cortex. To ease of understanding, we divide the swapping operation into five steps, similar to primitive operations underlying swapping in computer organization, while the circuit’s steps are not strictly non-overlapping in time.

1. **Memory holding in delay periods**. Two memory slots (*M*_1_ and *M*_2_) are in Up state to hold the WM items by receiving a sustained gain during the whole WM period, spanning from the Delay-1 to Delay-2 periods in the experiment (Fig. 1B). The need of sustained gain is analogous to the need of attention in loading and holding content in the WM [33]. In contrast, temporary slots (*T*_12_ and *T*_21_) are in Down state hence their activities are too low to hold any content during the Delay-1 period (Fig. 2E, light blue), even if they receive structured E inputs from memory slots (*M*_1_ and *M*_2_).
2. **Loading from memory slots to temporary slots**. The external sensory rule signal activates the swapping control circuit (Eq. 5; Fig. 3A, top purple line) that in turn increases the gain of temporary slots (*T*_12_ and *T*_21_). This switches on temporal slots into Up state through the cusp bifurcation (emerging a new fixed point with high activity; Fig. 2E, green dot), making them able to receive structured E inputs from memory slots (*T*_12_ *←M*_1_ and *T*_21_ *←M*_2_), corresponding to loading the content from memory slots. This corresponds to the step 1 in the swapping algorithm (Fig. 1C).
3. **“Erasing” the old content in memory slots**. The mutual inhibition between memory and temporary slots creates approximate winner-take-all activity between them (Eq. 4, Fig. 2B). When temporal slots build their responses aided by the gain from control circuits, they suppress memory slots’ responses into a very low level (Fig. 2F), effectively “erasing” the old content in memory slots^2^. Note that steps 2) and 3) happen concurrently and gradually.
4. **Loading from temporal slots to memory slots**. Although the swapping sensory rule is *transient*, the swapping control neuron self-sustain its response via recurrent excitation (Eq. 5, *w*_*c*_), and provides sustained gain to temporary slots. This enables temporary slots to reactivate subsequent memory slots and relay WM contents to them (Fig. 1C, step 2), completing the WM item swapping.
5. **Execution confirmation via inhibitory feedback from memory to control circuits**. The swapping command neuron receives inhibitory feedback from memory slots. After memory slots rebuild their responses, they will inhibit the control circuit and finally shut it down, as an internal confirmation showing that the swapping operation is executed. Without this internal feedback, the swapping control neurons will persistently activate temporary slots and repeat the swapping.

**Figure 3.**
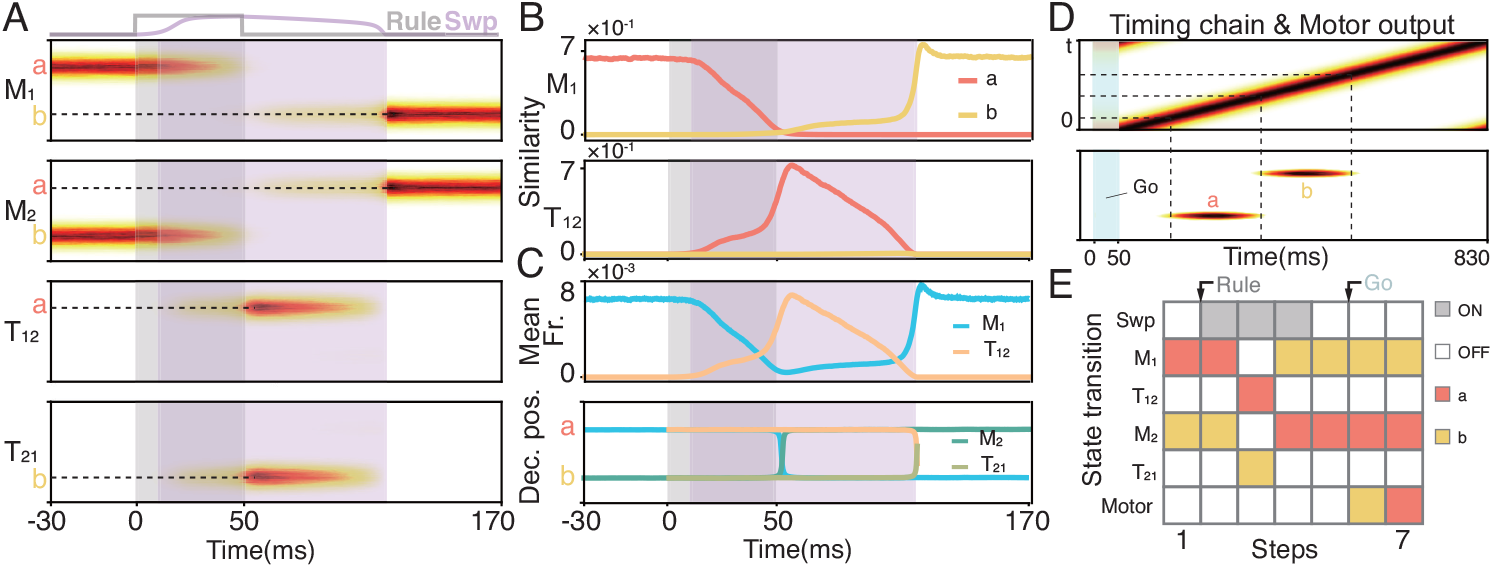
(A) Population activities of (temporary) memory slots. Shaded areas indicate the time windows when rule cue is presented (gray) and swapping control neuron fires (purple). (B) Similarity between neural activities of *M*_1_, *T*_12_ and presented stimulus. (C) Top: Mean firing rates of *M*_1_ and *T*_12_. Bottom: Decoded position from neural activities of 4 memory (temporary) slots. (D) Neural sequence internally generated in the output control sub-module (top) and the WM readout in the motor module (bottom). The WM items in memory slots are loaded into the motor module one by one dictated by the neural sequence in output control circuit. (E) The circuit activity with steps. White blocks: OFF states (low firing rate); gray or colored blocks: ON states (high firing rate).

Fig. 3A shows the joint activities of those memory and temporary slots and control circuits during the swapping operation. All the operations here are autonomous based on the recurrent dynamics.

## 3 The circuit for reading out working memory sequence

After the swapping operation, the animal receives a Go cue triggering motor responses to output the stored working memory sequence. We next attach another two components into the network: one is a output control sub-module which provides gain modulations on the memory slots to dictate them to run the program of motor readout, and another is a motor module for “generating” motor responses.

### A common motor readout region with threshold

There is a common motor region that reads out the content stored in memory slots in a fixed order. Note that the motor region corresponds to a different brain region from the memory slots recorded in the frontal cortex in the experiment. The motor region is also modeled as a CAN with the same internal structure as a memory slot and receives structured feedforward inputs from all memory slots (Eq. 2). The angular position encoded by the population activity in the motor region will directly trigger a movement along that direction, which was supported by the eye movement in superior colliculus for example [34].

Importantly, the motor readout region has a *threshold* to receive memory slots’ inputs, which may from the cortial basal ganglia loop [35]. During the WM period, the inputs from memory to the motor module cannot pass the motor threshold and hence not able to trigger motor outputs. This is realized by coordinating the feedforward weight from memory to the motor module with the motor threshold.

### The control circuits of motor readout: gain modulation on memory slots

There is another output control circuit providing gain modulations to the memory slots to implement WM sequence readout (Fig. 2B, top right), which is a separate sub-module from the swapping control circuit. In principle, we could use the output control circuit to increase the gain of memory slots one by one to pass the motor threshold to trigger the motor output. To realize this, the readout control circuit module is modeled as a widely used *timing sequence chain* of E neuron that internally generates a moving neural sequence along the chain [36–38] (see details in SI. S4). The neurons on the chain are topologically connected to memory slots in readout order, and then the moving sequence on the chain will enable the memory slots send content into the motor module sequentially (Fig. 4F).

**Figure 4.**
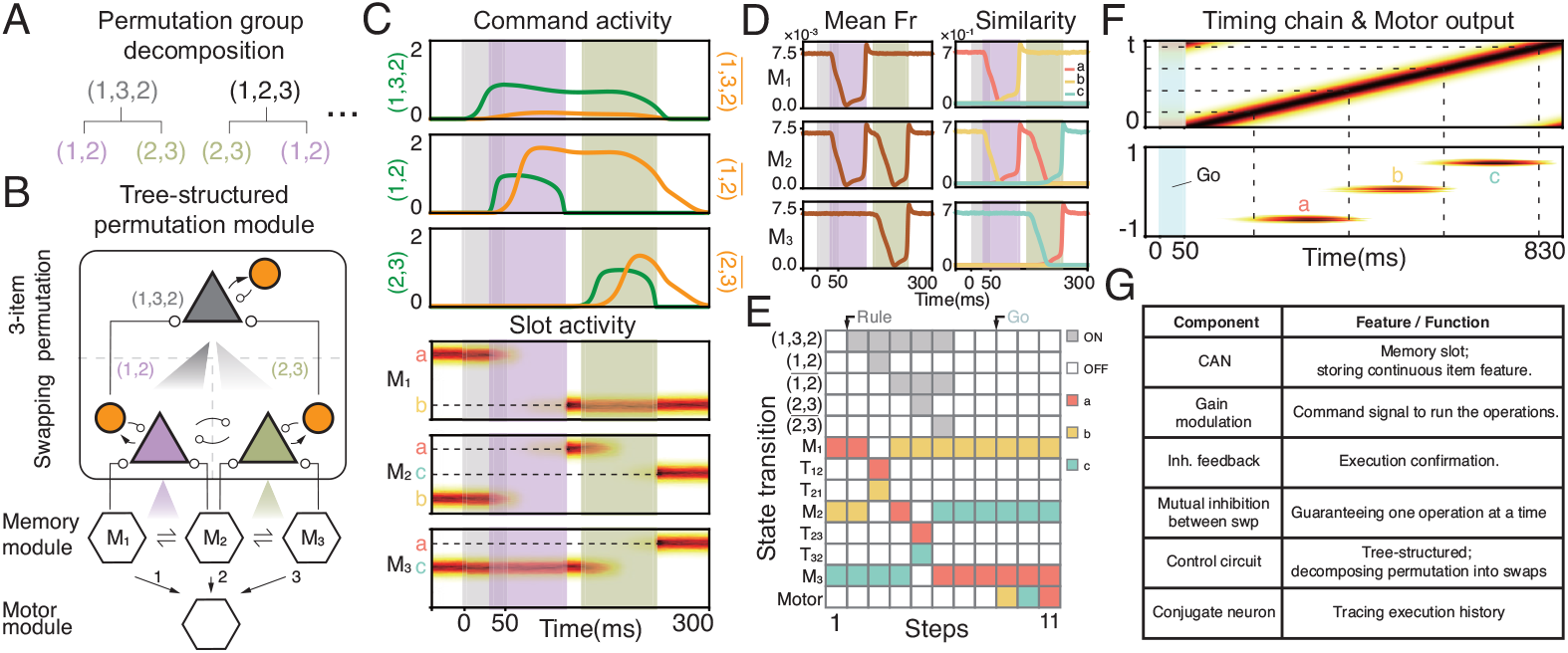
(A) Permutation group theory provides a strategy to decompose complex permutations into a swapping sequence. (B) A 3-item permutation can be implemented by executing two swappings. The gain input from the 3-permutation to swapping control neurons decreases with the order of swappings in the sequence. Triangle: permutation/swap control neurons; Orange circle: conjugate neurons; (C) Neural activities of control module and memory module. (D) Similarity and population firing rate in memory slots. (E) State transition. (F) Neural sequence of output control sub-module (top) and WM readout in motor module (bottom) (G) A table summarizing major components of the network.

## 4 Generalization to longer working memory sequence manipulation

Many cognitive tasks require storage and manipulation of longer WM sequences [39, 40], while the available data only has neuronal activities of two-item swapping [9]. Without direct evidence, we resort to normative math theories and combine them with the “designing principle” of the swapping circuit to explore a biologically plausible circuit for longer WM sequence manipulations.

### 4.1 Permutation group for swapping and reordering sequences

We use the algebraic group theory to formalize the two-item swapping operation and then generalize to more complicated sequence manipulations. Eventually, the group representation theory will provide insights into circuit architecture for multiple-item sequence manipulation as will be shown below. In theory, the two-item swapping forms a symmetric group *S*_2_. With the permutation cycle notation, the group *S*_2_ has two elements: () and (1, 2), where () is the identity operation that does not change the sequence, and (1, 2) denotes swapping the content in slot 1 and 2.

Similarly, all permutations of three-item sequences form the symmetric group *S*_3_ containing 6 permutations: {(),(1, 2),(2, 3),(1, 3),(1, 2, 3),(1, 3, 2)}, where the cycle notation (1, 2, 3) means moving slot 1’s original item to slot 2, slot 2’s original item to slot 3, and slot 3’s original item to slot 1, i.e., changing the original sequence *abc* to *cab*.

### 4.2 Permutation decomposition from group representation theory

The permutation group representation suggests a complex permutation can be decomposed into a series of simple swappings. For example, the permutation (1, 3, 2) can be decomposed into (Fig. 4A),

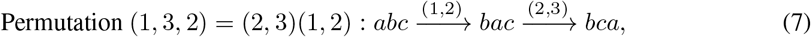

which means applying the permutation (1, 2) followed by the (2, 3), converting a three-item sequence

*abc* to *bca*. Table. S1 is the the group *S*_3_ multiplication table, which lists all possible decompositions.

## 5 The circuit for swapping 3-item working memory sequences

As a proof of concept, we develop a circuit model for 3-item permutations, i.e., *S*_3_ group, which can be easily generalized to manipulating longer WM sequences.

### 5.1 A tree-structured control circuit for permutation decomposition

Inspired by the decomposition of a complex permutation into a sequence of elementary swapping operations (Eq. 7), we propose a *tree-structured* control circuit module to implement complex permutations. In the control tree circuit, leaf nodes are binary swappings (Fig. 4B, triangles), each of which is the same as the one in previous swapping circuit (Fig. 2B, left) and provides gain modulations to memory modules. And the parents of the leaf nodes are the composition of swapping operations. In this way, the tree can be deepened to compose more complicated permutations, where the tree depth denotes the number of swappings in the composed permutation.

#### A parsimonious control tree

Although there are three swappings in *S*_3_, i.e., (1, 2),(2, 3),(1, 3), it is unnecessary to embed all of them in the leaf nodes. Instead, we only consider an example of swappings between *adjacent* memory slots, i.e., (1, 2) and (2, 3), which are sufficient to compose all *S*_3_ permutations (Table. S1). This implies that we only need four temporary slots in memory module: two between the memory slots *M*_1_ and *M*_2_, and another two between *M*_2_ and *M*_3_ (Fig. 4B). The adjacent swapping significantly reduces the number of swapping control neurons (tree’s leaf nodes) and temporary slots. For a general group *S*_*n*_ of *n*-item sequences, the adjacent swapping only requires *n −*1 swapping control neurons as the leaf nodes, while *S*_*n*_ group contains *n*(*n−* 1)*/*2 swappings. In principle, there are multiple choices of two swappings to compose *S*_3_, but we only consider one possible implementation in the present work.

#### Self-similarity for the interactions across tree hierarchy

Every node (permutation control neuron) in the control tree has similar dynamics with Eq. (5), and the interactions between a pair of nodes from two adjacent levels are similar to the interactions between swapping control neurons and memory slots in the previous swapping circuit (Fig. 4B). That is, the parent nodes (complex permutations) provide top-down gain modulations to the child nodes (simple permutation), and the child nodes provide bottom-up suppressive feedback to parent nodes to confirm the execution of simple permutation.

### 5.2 Generating swapping sequences by the control tree

An extra function required for the 3-item permutation circuit is to generate a sequence of swappings (Eq. 7), which is a different sequence from the WM sequence stored in the memory module. Hence, there are two sequences represented in the circuit. We next explore the mechanism of swapping sequence generation by the control circuit tree, which requires two additional mechanisms.

#### Spatial gradient of gain modulation determines the order of swappings

To generate a particular swapping sequence, the top-down modulations from the parent node on the tree should have a spatial gradient over the child nodes: parent node sends the largest gain to the first child node and the gain gradually decreases with the rank of the swapping within the swapping sequence (Fig. 4B), which enables that the earlier swappings will be activated first. Different parent nodes send different spatial gradients to the leaf nodes. The spatial gradient of gain is supported by a recent experiment [41] Moreover, to ensure that there is only one swapping being executed at a time, the two leaf nodes (swapping control neurons) mutually inhibit each other, forming a winner-take-all dynamics.

#### Conjugate neurons: register execution history and help generate swapping sequences

Each swapping control neuron is associated with a conjugate neuron (Fig. 4B, orange circles) that is driven by the control neuron and provides inhibitory feedback. The conjugate neuron will be activated once the corresponding swapping is executed, denoting the completion of the swapping. The conjugate neurons play two crucial roles: 1) tracing the execution history of swappings; 2) inhibiting the control neurons from returning back to executed operations (details in Sec. S3.3). When all swappings are executed, all conjugate neurons will suppress the top node (3-item permutation) to confirm the completion of the whole permutation.

### 5.3 The procedure of 3-item permutation in the circuit

We elaborate the procedures of 3-item permutation (1, 3, 2) in the circuit (Fig. 4B).

1. Activating the 3-item permutation node (1, 3, 2) with a transient external sensory rule signal.
2. The permutation node (1, 3, 2) modulates the gain of two leaf nodes (1, 2) and (2, 3) with spatial heterogeneity: larger gain to first swapping node (1, 2) and smaller to (2, 3).
3. The first node (1, 2) is ON and triggers swapping between memory slots *M*_1_ and *M*_2_ (the same as Sec. 2.4).
4. Memory slots (*M*_1_, *M*_2_) switch off node (1, 2) and meanwhile its conjugate 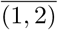 is turned ON representing the execution of (1, 2).
5. The next swapping node (2, 3) is activated since the node (1, 2) is OFF (release from mutual inhibition), triggering the second swapping between *M*_2_ and *M*_3_
6. When all conjugate neurons at the root level are ON, their inhibitory feedback in total will be sufficient to shut down the parent node (1, 3, 2) to confirm the execution of the whole permutation.

In addition, the 3-item permutation circuit is automatically downward compatible to perform single swappings, when the external rule signal is directly fed into the leaf node.

## 6 Conclusion and Discussion

The WM manipulation is central to cognitive reasoning while its circuit mechanism remains elusive. We propose a recurrent neural circuit model for WM sequence manipulation, including two-item swapping and 3-item permutations, and reproduce key observations in a recent experiment [9]. The circuit has a modular architecture consisting of memory module and tree-structured control module, each of which is based on canonical circuit motifs and operations, including CAN-based memory slots, gain modulation, and mutual inhibition. It provides several profound implications (Fig. 4G):

1. The simple gain modulation can orchestrate complex programs in memory modules.
2. The tree-structured control circuit, directly specified from the permutation group theory, can decompose a complex permutation into compositions of elementary swaps.
3. The inhibitory feedback from memory to control module can confirm program execution.
4. The mutual inhibition between elementary swaps ensure only one operation is ran at a time. Considering the swap and permutation are elementary operations, the proposed circuit model helps us to understand the circuit mechanism of cognitive reasoning and inspire new AI reasoning systems.

### Experimental support of the circuit motifs and operations in the model

1. CAN-based memory slots: supported by extensive experimental and modeling studies [10–14].
2. Control module. Although lacking direct neurophysiological experimental evidence, it is broadly consistent with the idea of selective gating in conceptual models of working memory [42, 43, 41] which relied on spatial or temporal gating of signals for information encoding and retrieval.
3. Tree-structure of the control module. It is directly supported by a recent WM sequence experiment [44]. In principle, the tree structure can be generalized to the hyperbolic geometry (a tree with continuous layers) which was discovered in the brain, e.g., olfactory system [45] and hippocampus [46], and is widely used in ML to represent hierarchical structures, e.g., word embeddings [47].
4. Decomposition of complex operations. This is widely observed across a wide range of neural systems, e.g., motor systems [48–50], prefrontal cortex [51], language [52], visual systems [53].
5. Gain modulation: widely observed in the brain, e.g., attention [22], memory [23], etc [24].
6. Inhibitory feedback for execution confirmation. Using inhibition to confirm is consistent with the *inhibition of return* in spatial attention that prevents attention returning to an earlier location [54].

#### The neural circuit mechanism of symbol

How to emerge and represent discrete symbols in continuous valued neural circuits is a fundamental question in neuroscience and AI. The symbol formation requires a boundary to separate the WM content. In our circuit model, a symbol corresponds to a WM item, e.g., a/b/c in Fig. 1A, and is represented by a specialized memory slot of continuous attractor dynamics, which was also used in a recent study [55]. And the continuous feature associated with a symbol (e.g., Fig. 1B, angular position) is represented by the bump position in the attractor dynamics. Importantly, the separation between symbols corresponds to the separation (boundary) between memory slots, which can be determined by the range of inhibitory connections (Fig. 2C, red): The E neurons within a memory slot share their own inhibitory neuron pool (Eq. 3, denominator). This implies inhibitory neurons might be crucial for forming symbols in neural circuits.

#### Comparison to other work

To our best knowledge, the present study is the first experiment-supported neural circuit model of WM sequence manipulation, since the experiment [9] is the first to show the neural substrate of WM swapping. Compared to programming network models in ML (e.g., [56, 6]), most of them use *gating* to directly change the network weight (matrix), while, our circuit model relies on gain modulation to uniformly change neuronal activation functions within a population (slot or control sub-module). This gain control requires less spatial resolution than the weight gating in artificial networks, and may be a biologically plausible way to improve ML models.

#### Generalization to complex programs

The modular circuit architecture empowers the proposed circuit model to easily generalize to complex programs by simple control module upgrade, without changing memory slots. For example, as illustrated by the swapping to 3-item permutations, we only need to increase the height of the control circuit tree. Considering that the permutation is an elementary operation in programming, the permutation control tree can be used as a basis to be called by a higher control node representing a complex program. In general, it implies that running a new program only needs to learn a control sequence within the control circuit, minimizing the modification of the circuit. Furthermore, it implies that different programs correspond to different sequences within the circuit, consistent with the concept of spatial computing in the cortex [57, 58].

#### Limitation and future work

Our work doesn’t include the process of loading external sensory inputs into memory slots, which also needs a control coordination to determine the routing from sensory inputs to memory slots [33]. Limited by the space, we only consider permuting a 3-item WM sequences. Generalizing to a longer WM sequences manipulation requires a higher control circuit tree, whose neuronal number may grow with the order of 𝒪 (2^*h*^) with *h* the number of tree layers. Nevertheless, the high demand for control circuit neurons may be partially alleviated by the limited capacity of the WM system in the brain, i.e., about 4-7 items [39, 40]. These form our future work.

## Acknowledgments

W.H.Z. is supported by the UT Southwestern Medical Center Endowed Scholars program. S.W. is supported by the Science Fund for Creative Research Groups of the National Natural Science Foundation of China (https://www.nsfc.gov.cn, T2421004) and the National Key Research and Development Program of China (2024YFF1206500).

## NeurIPS Paper Checklist

### 1. Claims

Question: Do the main claims made in the abstract and introduction accurately reflect the paper’s contributions and scope?

Answer: [Yes]

Justification: We summarize our claims and contribution clearly in the Abstract and Introduction, and also discuss the limitation and extension of the current work in Discussion.

Guidelines:

- The answer NA means that the abstract and introduction do not include the claims made in the paper.
- The abstract and/or introduction should clearly state the claims made, including the contributions made in the paper and important assumptions and limitations. A No or NA answer to this question will not be perceived well by the reviewers.
- The claims made should match theoretical and experimental results, and reflect how much the results can be expected to generalize to other settings.
- It is fine to include aspirational goals as motivation as long as it is clear that these goals are not attained by the paper.

### 2. Limitations

Question: Does the paper discuss the limitations of the work performed by the authors? Answer: [Yes]

Justification: We discussed the limitations of the present work in the Discussion section. Guidelines:

- The answer NA means that the paper has no limitation while the answer No means that the paper has limitations, but those are not discussed in the paper.
- The authors are encouraged to create a separate “Limitations” section in their paper.
- The paper should point out any strong assumptions and how robust the results are to violations of these assumptions (e.g., independence assumptions, noiseless settings, model well-specification, asymptotic approximations only holding locally). The authors should reflect on how these assumptions might be violated in practice and what the implications would be.
- The authors should reflect on the scope of the claims made, e.g., if the approach was only tested on a few datasets or with a few runs. In general, empirical results often depend on implicit assumptions, which should be articulated.
- The authors should reflect on the factors that influence the performance of the approach. For example, a facial recognition algorithm may perform poorly when image resolution is low or images are taken in low lighting. Or a speech-to-text system might not be used reliably to provide closed captions for online lectures because it fails to handle technical jargon.
- The authors should discuss the computational efficiency of the proposed algorithms and how they scale with dataset size.
- If applicable, the authors should discuss possible limitations of their approach to address problems of privacy and fairness.
- While the authors might fear that complete honesty about limitations might be used by reviewers as grounds for rejection, a worse outcome might be that reviewers discover limitations that aren’t acknowledged in the paper. The authors should use their best judgment and recognize that individual actions in favor of transparency play an important role in developing norms that preserve the integrity of the community. Reviewers will be specifically instructed to not penalize honesty concerning limitations.

### 3. Theory assumptions and proofs

Question: For each theoretical result, does the paper provide the full set of assumptions and a complete (and correct) proof?

Answer: [Yes]

Justification: We clearly provided a full set of assumptions and proof in the main and supplementary texts.

Guidelines:

- The answer NA means that the paper does not include theoretical results.
- All the theorems, formulas, and proofs in the paper should be numbered and cross-referenced.
- All assumptions should be clearly stated or referenced in the statement of any theorems.
- The proofs can either appear in the main paper or the supplemental material, but if they appear in the supplemental material, the authors are encouraged to provide a short proof sketch to provide intuition.
- Inversely, any informal proof provided in the core of the paper should be complemented by formal proofs provided in appendix or supplemental material.
- Theorems and Lemmas that the proof relies upon should be properly referenced.

### 4. Experimental result reproducibility

Question: Does the paper fully disclose all the information needed to reproduce the main experimental results of the paper to the extent that it affects the main claims and/or conclusions of the paper (regardless of whether the code and data are provided or not)?

Answer: [Yes]

Justification: We provided a table of all parameters and devices in the Supplementary Information.

Guidelines:

- The answer NA means that the paper does not include experiments.
- If the paper includes experiments, a No answer to this question will not be perceived well by the reviewers: Making the paper reproducible is important, regardless of whether the code and data are provided or not.
- If the contribution is a dataset and/or model, the authors should describe the steps taken to make their results reproducible or verifiable.
- Depending on the contribution, reproducibility can be accomplished in various ways. For example, if the contribution is a novel architecture, describing the architecture fully might suffice, or if the contribution is a specific model and empirical evaluation, it may be necessary to either make it possible for others to replicate the model with the same dataset, or provide access to the model. In general. releasing code and data is often one good way to accomplish this, but reproducibility can also be provided via detailed instructions for how to replicate the results, access to a hosted model (e.g., in the case of a large language model), releasing of a model checkpoint, or other means that are appropriate to the research performed.
- While NeurIPS does not require releasing code, the conference does require all submissions to provide some reasonable avenue for reproducibility, which may depend on the nature of the contribution. For example
  a. If the contribution is primarily a new algorithm, the paper should make it clear how to reproduce that algorithm.
  b. If the contribution is primarily a new model architecture, the paper should describe the architecture clearly and fully.
  c. If the contribution is a new model (e.g., a large language model), then there should either be a way to access this model for reproducing the results or a way to reproduce the model (e.g., with an open-source dataset or instructions for how to construct the dataset).
  d. We recognize that reproducibility may be tricky in some cases, in which case authors are welcome to describe the particular way they provide for reproducibility. In the case of closed-source models, it may be that access to the model is limited in some way (e.g., to registered users), but it should be possible for other researchers to have some path to reproducing or verifying the results.

### 5. Open access to data and code

Question: Does the paper provide open access to the data and code, with sufficient instructions to faithfully reproduce the main experimental results, as described in supplemental material?

Answer: [Yes]

Justification: We included our codes in the Supplementary Information. Guidelines:

- The answer NA means that paper does not include experiments requiring code.
- Please see the NeurIPS code and data submission guidelines (https://nips.cc/public/guides/CodeSubmissionPolicy) for more details.
- While we encourage the release of code and data, we understand that this might not be possible, so “No” is an acceptable answer. Papers cannot be rejected simply for not including code, unless this is central to the contribution (e.g., for a new open-source benchmark).
- The instructions should contain the exact command and environment needed to run to reproduce the results. See the NeurIPS code and data submission guidelines (https://nips.cc/public/guides/CodeSubmissionPolicy) for more details.
- The authors should provide instructions on data access and preparation, including how to access the raw data, preprocessed data, intermediate data, and generated data, etc.
- The authors should provide scripts to reproduce all experimental results for the new proposed method and baselines. If only a subset of experiments are reproducible, they should state which ones are omitted from the script and why.
- At submission time, to preserve anonymity, the authors should release anonymized versions (if applicable).
- Providing as much information as possible in supplemental material (appended to the paper) is recommended, but including URLs to data and code is permitted.

### 6. Experimental setting/details

Question: Does the paper specify all the training and test details (e.g., data splits, hyperparameters, how they were chosen, type of optimizer, etc.) necessary to understand the results?

Answer: [Yes]

Justification: While this work is not associated with model training and testing, the details of model simulation is presented in Sec. S5.

Guidelines:

- The answer NA means that the paper does not include experiments.
- The experimental setting should be presented in the core of the paper to a level of detail that is necessary to appreciate the results and make sense of them.
- The full details can be provided either with the code, in appendix, or as supplemental material.

### 7. Experiment statistical significance

Question: Does the paper report error bars suitably and correctly defined or other appropriate information about the statistical significance of the experiments?

Answer: [NA]

Justification: This work is a theoretical neuroscience study that only involves group theory and dynamical systems but does not involve statistics.

Guidelines:

- The answer NA means that the paper does not include experiments.
- The authors should answer “Yes” if the results are accompanied by error bars, confidence intervals, or statistical significance tests, at least for the experiments that support the main claims of the paper.
- The factors of variability that the error bars are capturing should be clearly stated (for example, train/test split, initialization, random drawing of some parameter, or overall run with given experimental conditions).
- The method for calculating the error bars should be explained (closed form formula, call to a library function, bootstrap, etc.)
- The assumptions made should be given (e.g., Normally distributed errors).
- It should be clear whether the error bar is the standard deviation or the standard error of the mean.
- It is OK to report 1-sigma error bars, but one should state it. The authors should preferably report a 2-sigma error bar than state that they have a 96% CI, if the hypothesis of Normality of errors is not verified.
- For asymmetric distributions, the authors should be careful not to show in tables or figures symmetric error bars that would yield results that are out of range (e.g. negative error rates).
- If error bars are reported in tables or plots, The authors should explain in the text how they were calculated and reference the corresponding figures or tables in the text.

### 8. Experiments compute resources

Question: For each experiment, does the paper provide sufficient information on the computer resources (type of compute workers, memory, time of execution) needed to reproduce the experiments?

Answer: [Yes]

Justification: Details can be found in Sec. S5. Guidelines:

- The answer NA means that the paper does not include experiments.
- The paper should indicate the type of compute workers CPU or GPU, internal cluster, or cloud provider, including relevant memory and storage.
- The paper should provide the amount of compute required for each of the individual experimental runs as well as estimate the total compute.
- The paper should disclose whether the full research project required more compute than the experiments reported in the paper (e.g., preliminary or failed experiments that didn’t make it into the paper).

### 9. Code of ethics

Question: Does the research conducted in the paper conform, in every respect, with the NeurIPS Code of Ethics https://neurips.cc/public/EthicsGuidelines?

Answer: [Yes]

Justification: We confirm our study conform the Code of Ethics. Guidelines:

- The answer NA means that the authors have not reviewed the NeurIPS Code of Ethics.
- If the authors answer No, they should explain the special circumstances that require a deviation from the Code of Ethics.
- The authors should make sure to preserve anonymity (e.g., if there is a special consideration due to laws or regulations in their jurisdiction).

### 10. Broader impacts

Question: Does the paper discuss both potential positive societal impacts and negative societal impacts of the work performed?

Answer: [NA]

Justification: This is a theoretical study for basic science research and will not have direct societal impacts.

Guidelines:

- The answer NA means that there is no societal impact of the work performed.
- If the authors answer NA or No, they should explain why their work has no societal impact or why the paper does not address societal impact.
- Examples of negative societal impacts include potential malicious or unintended uses (e.g., disinformation, generating fake profiles, surveillance), fairness considerations (e.g., deployment of technologies that could make decisions that unfairly impact specific groups), privacy considerations, and security considerations.
- The conference expects that many papers will be foundational research and not tied to particular applications, let alone deployments. However, if there is a direct path to any negative applications, the authors should point it out. For example, it is legitimate to point out that an improvement in the quality of generative models could be used to generate deepfakes for disinformation. On the other hand, it is not needed to point out that a generic algorithm for optimizing neural networks could enable people to train models that generate Deepfakes faster.
- The authors should consider possible harms that could arise when the technology is being used as intended and functioning correctly, harms that could arise when the technology is being used as intended but gives incorrect results, and harms following from (intentional or unintentional) misuse of the technology.
- If there are negative societal impacts, the authors could also discuss possible mitigation strategies (e.g., gated release of models, providing defenses in addition to attacks, mechanisms for monitoring misuse, mechanisms to monitor how a system learns from feedback over time, improving the efficiency and accessibility of ML).

### 11. Safeguards

Question: Does the paper describe safeguards that have been put in place for responsible release of data or models that have a high risk for misuse (e.g., pretrained language models, image generators, or scraped datasets)?

Answer: [NA]

Justification: This is a theoretical study for basic science research and will not incur any risk.

Guidelines:

- The answer NA means that the paper poses no such risks.
- Released models that have a high risk for misuse or dual-use should be released with necessary safeguards to allow for controlled use of the model, for example by requiring that users adhere to usage guidelines or restrictions to access the model or implementing safety filters.
- Datasets that have been scraped from the Internet could pose safety risks. The authors should describe how they avoided releasing unsafe images.
- We recognize that providing effective safeguards is challenging, and many papers do not require this, but we encourage authors to take this into account and make a best faith effort.

### 12. Licenses for existing assets

Question: Are the creators or original owners of assets (e.g., code, data, models), used in the paper, properly credited and are the license and terms of use explicitly mentioned and properly respected?

Answer: [Yes]

Justification: We adapted a figure from an experimental paper (Fig. 1AB) and acknowledged that in the text.

Guidelines:

- The answer NA means that the paper does not use existing assets.
- The authors should cite the original paper that produced the code package or dataset.
- The authors should state which version of the asset is used and, if possible, include a URL.
- The name of the license (e.g., CC-BY 4.0) should be included for each asset.
- For scraped data from a particular source (e.g., website), the copyright and terms of service of that source should be provided.
- If assets are released, the license, copyright information, and terms of use in the package should be provided. For popular datasets, paperswithcode.com/datasets has curated licenses for some datasets. Their licensing guide can help determine the license of a dataset.
- For existing datasets that are re-packaged, both the original license and the license of the derived asset (if it has changed) should be provided.
- If this information is not available online, the authors are encouraged to reach out to the asset’s creators.

### 13. New assets

Question: Are new assets introduced in the paper well documented and is the documentation provided alongside the assets?

Answer: [Yes]

Justification: And the code of simulating the model is uploaded. Guidelines:

- The answer NA means that the paper does not release new assets.
- Researchers should communicate the details of the dataset/code/model as part of their submissions via structured templates. This includes details about training, license, limitations, etc.
- The paper should discuss whether and how consent was obtained from people whose asset is used.
- At submission time, remember to anonymize your assets (if applicable). You can either create an anonymized URL or include an anonymized zip file.

### 14. Crowdsourcing and research with human subjects

Question: For crowdsourcing experiments and research with human subjects, does the paper include the full text of instructions given to participants and screenshots, if applicable, as well as details about compensation (if any)?

Answer: [NA]

Justification: This paper does not involve crowdsourcing nor research with human subjects. Guidelines:

- The answer NA means that the paper does not involve crowdsourcing nor research with human subjects.
- Including this information in the supplemental material is fine, but if the main contribution of the paper involves human subjects, then as much detail as possible should be included in the main paper.
- According to the NeurIPS Code of Ethics, workers involved in data collection, curation, or other labor should be paid at least the minimum wage in the country of the data collector.

### 15. Institutional review board (IRB) approvals or equivalent for research with human subjects

Question: Does the paper describe potential risks incurred by study participants, whether such risks were disclosed to the subjects, and whether Institutional Review Board (IRB) approvals (or an equivalent approval/review based on the requirements of your country or institution) were obtained?

Answer: [NA]

Justification: This paper does not involve crowdsourcing nor research with human subjects. Guidelines:

- The answer NA means that the paper does not involve crowdsourcing nor research with human subjects.
- Depending on the country in which research is conducted, IRB approval (or equivalent) may be required for any human subjects research. If you obtained IRB approval, you should clearly state this in the paper.
- We recognize that the procedures for this may vary significantly between institutions and locations, and we expect authors to adhere to the NeurIPS Code of Ethics and the guidelines for their institution.
- For initial submissions, do not include any information that would break anonymity (if applicable), such as the institution conducting the review.

### 16. Declaration of LLM usage

Question: Does the paper describe the usage of LLMs if it is an important, original, or non-standard component of the core methods in this research? Note that if the LLM is used only for writing, editing, or formatting purposes and does not impact the core methodology, scientific rigorousness, or originality of the research, declaration is not required.

Answer: [NA]

Justification: The core method development in this research does not involve LLMs as any important, original, or non-standard components.

Guidelines:

- The answer NA means that the core method development in this research does not involve LLMs as any important, original, or non-standard components.
- Please refer to our LLM policy (https://neurips.cc/Conferences/2025/LLM) for what should or should not be described.

## Footnotes

2 This is not completely erase, but the low response of memory slots make them easier to load new content.

## References

[1] Philip Nicholas Johnson-Laird. Mental models: Towards a cognitive science of language, inference, and consciousness. Number 6. Harvard University Press, 1983.

[2] A. M. Turing. Computing machinery and intelligence. Mind, LIX(236):433–460, 10 1950.

[3] Joaquin M Fuster and Garrett E Alexander. Neuron activity related to short-term memory. Science, 173(3997):652–654, 1971.

[4] Earl K Miller and Jonathan D Cohen. An integrative theory of prefrontal cortex function. Annual review of neuroscience, 24(1):167–202, 2001.

[5] Ashish Vaswani, Noam Shazeer, Niki Parmar, Jakob Uszkoreit, Llion Jones, Aidan N Gomez, Łukasz Kaiser, and Illia Polosukhin. Attention is all you need. Advances in neural information processing systems, 30, 2017.

[6] Alex Graves, Greg Wayne, Malcolm Reynolds, Tim Harley, Ivo Danihelka, Agnieszka Grabska-Barwińska, Sergio Gómez Colmenarejo, Edward Grefenstette, Tiago Ramalho, John Agapiou, et al. Hybrid computing using a neural network with dynamic external memory. Nature, 538(7626):471–476, 2016.

[7] Alec Radford, Karthik Narasimhan, Tim Salimans, Ilya Sutskever, et al. Improving language understanding by generative pre-training. 2018.

[8] Jacob Devlin, Ming-Wei Chang, Kenton Lee, and Kristina Toutanova. Bert: Pre-training of deep bidirectional transformers for language understanding. In Proceedings of the 2019 conference of the North American chapter of the association for computational linguistics: human language technologies, volume 1 (long and short papers), pages 4171–4186, 2019.

[9] Zhenghe Tian, Jingwen Chen, Cong Zhang, Bin Min, Bo Xu, and Liping Wang. Mental programming of spatial sequences in working memory in the macaque frontal cortex. Science, 385(6716):eadp6091, 2024.

[10] Albert Compte, Christos Constantinidis, Jesper Tegnér, Sridhar Raghavachari, Matthew V Chafee, Patricia S Goldman-Rakic, and Xiao-Jing Wang. Temporally irregular mnemonic persistent activity in prefrontal neurons of monkeys during a delayed response task. Journal of neurophysiology, 90(5):3441–3454, 2003.

[11] Dante Francisco Wasmuht, Eelke Spaak, Timothy J Buschman, Earl K Miller, and Mark G Stokes. Intrinsic neuronal dynamics predict distinct functional roles during working memory. Nature communications, 9(1):3499, 2018.

[12] Klaus Wimmer, Duane Q Nykamp, Christos Constantinidis, and Albert Compte. Bump attractor dynamics in prefrontal cortex explains behavioral precision in spatial working memory. Nature neuroscience, 17(3):431, 2014.

[13] Albert Compte, Nicolas Brunel, Patricia S Goldman-Rakic, and Xiao-Jing Wang. Synaptic mechanisms and network dynamics underlying spatial working memory in a cortical network model. Cerebral cortex, 10(9):910–923, 2000.

[14] X-J Wang, Jesper Tegnér, C Constantinidis, and Patricia S Goldman-Rakic. Division of labor among distinct subtypes of inhibitory neurons in a cortical microcircuit of working memory. Proceedings of the National Academy of Sciences, 101(5):1368–1373, 2004.

[15] Si Wu, Hiroyuki Nakahara, and Shun-Ichi Amari. Population coding with correlation and an unfaithful model. Neural Computation, 13(4):775–797, 2001.

[16] Si Wu, Shun-ichi Amari, and Hiroyuki Nakahara. Population coding and decoding in a neural field: a computational study. Neural Computation, 14(5):999–1026, 2002.

[17] Si Wu, Kosuke Hamaguchi, and Shun-ichi Amari. Dynamics and computation of continuous attractors. Neural Computation, 20(4):994–1025, 2008.

[18] MA Stephens. Random walk on a circle. Biometrika, 50(3/4):385–390, 1963.

[19] Michael J Tarr and Steven Pinker. Mental rotation and orientation-dependence in shape recognition. Cognitive psychology, 21(2):233–282, 1989.

[20] Earl K Miller, Mikael Lundqvist, and André M Bastos. Working memory 2.0. Neuron, 100(2):463–475, 2018.

[21] John D Dixon and Brian Mortimer. Permutation groups, volume 163. Springer Science & Business Media, 1996.

[22] Douglas A Ruff and Marlene R Cohen. Simultaneous multi-area recordings suggest that attention improves performance by reshaping stimulus representations. Nature neuroscience, 22(10):1669–1676, 2019.

[23] Sean M Polyn, Kenneth A Norman, and Michael J Kahana. A context maintenance and retrieval model of organizational processes in free recall. Psychological review, 116(1):129, 2009.

[24] Ramanujan Srinath, Martyna M Czarnik, and Marlene R Cohen. Coordinated response modulations enable flexible use of visual information. bioRxiv, 2024.

[25] R Ben-Yishai, R Lev Bar-Or, and H Sompolinsky. Theory of orientation tuning in visual cortex. Proceedings of the National Academy of Sciences, 92(9):3844–3848, 1995.

[26] James J Knierim and Kechen Zhang. Attractor dynamics of spatially correlated neural activity in the limbic system. Annual review of neuroscience, 35:267–285, 2012.

[27] Si Wu, KY Michael Wong, CC Alan Fung, Yuanyuan Mi, and Wenhao Zhang. Continuous attractor neural networks: candidate of a canonical model for neural information representation. F1000Research, 5, 2016.

[28] Mikail Khona and Ila R. Fiete. Attractor and integrator networks in the brain. Nature Reviews Neuroscience, 23(12):744–766, December 2022.

[29] Sophie Deneve, Peter E Latham, and Alexandre Pouget. Reading population codes: a neural implementation of ideal observers. Nature Neuroscience, 2(8):740–745, 1999.

[30] Matteo Carandini and David J Heeger. Normalization as a canonical neural computation. Nature Reviews Neuroscience, 13(1):51–62, 2012.

[31] Cristopher M Niell. Cell types, circuits, and receptive fields in the mouse visual cortex. Annual review of neuroscience, 38:413–431, 2015.

[32] John Guckenheimer and Yuri A Kuznetsov. Cusp bifurcation. Scholarpedia, 2(4):1852, 2007.

[33] Jingwen Chen, Cong Zhang, Peiyao Hu, Bin Min, and Liping Wang. Flexible control of sequence working memory in the macaque frontal cortex. Neuron, 112(20):3502–3514, 2024.

[34] Choongkil Lee, William H. Rohrer, and David L. Sparks. Population coding of saccadic eye movements by neurons in the superior colliculus. Nature, 332(6162):357–360, 1988.

[35] R Kaji. Basal ganglia as a sensory gating devise for motor control. The journal of medical investigation: JMI, 48(3-4):142–146, 2001.

[36] Kanaka Rajan, Christopher D Harvey, and David W Tank. Recurrent network models of sequence generation and memory. Neuron, 90(1):128–142, 2016.

[37] Manuel Beiran, Nicolas Meirhaeghe, Hansem Sohn, Mehrdad Jazayeri, and Srdjan Ostojic. Parametric control of flexible timing through low-dimensional neural manifolds. Neuron, 111(5):739–753, 2023.

[38] Junfeng Zuo, Xiao Liu, Ying Nian Wu, Si Wu, and Wenhao Zhang. A recurrent neural circuit mechanism of temporal-scaling equivariant representation. Advances in Neural Information Processing Systems, 36, 2023.

[39] Steven J Luck and Edward K Vogel. The capacity of visual working memory for features and conjunctions. Nature, 390(6657):279–281, 1997.

[40] Nelson Cowan. The magical number 4 in short-term memory: A reconsideration of mental storage capacity. Behavioral and brain sciences, 24(1):87–114, 2001.

[41] Edward Awh and Edward K Vogel. Working memory needs pointers. Trends in Cognitive Sciences, 2024.

[42] Sridhar Raghavachari, Michael J Kahana, Daniel S Rizzuto, Jeremy B Caplan, Matthew P Kirschen, Blaise Bourgeois, Joseph R Madsen, and John E Lisman. Gating of human theta oscillations by a working memory task. Journal of Neuroscience, 21(9):3175–3183, 2001.

[43] Timothy J Buschman and Earl K Miller. Working memory is complex and dynamic, like your thoughts. Journal of cognitive neuroscience, 35(1):17–23, 2022.

[44] Ying Fan, Muzhi Wang, Fang Fang, Nai Ding, and Huan Luo. Two-dimensional neural geometry underpins hierarchical organization of sequence in human working memory. Nature Human Behaviour, 9(2):360–375, 2025.

[45] Yuansheng Zhou, Brian H Smith, and Tatyana O Sharpee. Hyperbolic geometry of the olfactory space. Science advances, 4(8):eaaq1458, 2018.

[46] Huanqiu Zhang, P Dylan Rich, Albert K Lee, and Tatyana O Sharpee. Hippocampal spatial representations exhibit a hyperbolic geometry that expands with experience. Nature Neuroscience, 26(1):131–139, 2023.

[47] Maximillian Nickel and Douwe Kiela. Poincaré embeddings for learning hierarchical representations. Advances in neural information processing systems, 30, 2017.

[48] Ferdinando A Mussa-Ivaldi and Emilio Bizzi. Motor learning through the combination of primitives. Philosophical Transactions of the Royal Society of London. Series B: Biological Sciences, 355(1404):1755–1769, 2000.

[49] Emanuel Todorov. Compositionality of optimal control laws. Advances in neural information processing systems, 22, 2009.

[50] Robert C Berwick, Kazuo Okanoya, Gabriel JL Beckers, and Johan J Bolhuis. Songs to syntax: the linguistics of birdsong. Trends in cognitive sciences, 15(3):113–121, 2011.

[51] Feng-Kuei Chiang and Joni D Wallis. Neuronal encoding in prefrontal cortex during hierarchical reinforcement learning. Journal of Cognitive Neuroscience, 30(8):1197–1208, 2018.

[52] Steven M Frankland and Joshua D Greene. Concepts and compositionality: in search of the brain’s language of thought. Annual review of psychology, 71(1):273–303, 2020.

[53] Tai Sing Lee. The visual system’s internal model of the world. Proceedings of the IEEE, 103(8):1359–1378, 2015.

[54] Raymond M Klein. Inhibition of return. Trends in cognitive sciences, 4(4):138–147, 2000.

[55] Andrew Nam, Eric Elmoznino, Nikolay Malkin, James McClelland, Yoshua Bengio, and Guillaume Lajoie. Discrete, compositional, and symbolic representations through attractor dynamics. arXiv preprint 2310.01807, 2023.

[56] Alex Graves, Greg Wayne, and Ivo Danihelka. Neural turing machines. arXiv preprint 1410.5401, 2014.

[57] Mikael Lundqvist, Scott L Brincat, Jonas Rose, Melissa R Warden, Timothy J Buschman, Earl K Miller, and Pawel Herman. Working memory control dynamics follow principles of spatial computing. Nature Communications, 14(1):1429, 2023.

[58] Mikael Lundqvist, Earl K Miller, Jonatan Nordmark, Johan Liljefors, and Pawel Herman. Beta: bursts of cognition. Trends in cognitive sciences, 2024.

