## Supplemental Information for "Towards a mental programming neural circuit: Insights from working memory sequence manipulation"

**Junfeng Zuo[1,3], Cheng Xue[2], Si Wu[1], Wen-Hao Zhang[3,4] ***
[1]Peking-Tsinghua Center for Life Sciences, Academy for Advanced Interdisciplinary Studies, School of Psychology and Cognitive Sciences, IDG/McGovern Institute for Brain Research, Center of Quantitative Biology, Peking University.
[2]Department of Neurobiology, University of Chicago.
[3]Lyda Hill Department of Bioinformatics, UT Southwestern Medical Center.
[4]O'Donnell Brain Institute, UT Southwestern Medical Center.

**S4 Motor readout** **7**

**S5 Simulation details** **7**

**S6 Supplementary figures** **7**

Preprint. Under review.

#### S1 Slot representation

Without losing generality, we assume that each stimulus can be represented by a 1-dimensional variable, and that every stimulus is sampled from a common space, such as the direction space $[-\pi, \pi)$ (Fig. 1). Therefore, it is intuitively straightforward to model each slot as a 1-dimensional continuous attractor network (CAN), which can hold the memory of the stimulus in persistent neural activities.

##### S1.1 Continuous attractor network

The dynamics of a 1D-CAN can be written as:

$$\tau \dot{u}^E(\theta, t) = -u^E(\theta, t) + \rho \mathsf{W}_{\mathsf{rec}}(\theta) \star r^E(\theta, t) + I, \tag{S1a}$$

$$\tau_i \dot{r}^I(t) = -r^I(t) + \rho w^{IE} \int [u^E(\theta', t)]_+^2 \mathsf{d}\theta' \tag{S1b}$$

$$r^E(\theta, t) = \frac{g(t) \cdot [u^E(\theta, t)]_+^2}{1 + k \cdot r^I(t)}, \tag{S1c}$$

where $u(\theta, t), r(\theta, t)$ and $r^I(t)$ represent synaptic input, firing rate of excitatory neurons and firing rate of interneurons. $\tau, \rho, k$ are time constant, neuron density and inhibition strength respectively. $\theta$ represents the preferred stimulus of the corresponding excitatory neurons.$[\cdot]_+$ denotes the ReLU function where $[x]_+ = \mathsf{max}(x, 0)$. $I$ is external input, combining sensory input from the environment and projection from other slots. The inter-projection between slots can be described by $\mathsf{W}_{\mathsf{proj}}(\theta) \star r^E(\theta, t)$.

The excitatory recurrent and inter connections within slots can be written as Gaussian-shaped functions $\mathsf{W}_{\mathsf{x}}(\theta)$, which can be described by:

$$\mathsf{W}_{\mathsf{x}}(\theta) = w_{\mathsf{x}}^{EE} \cdot (\sqrt{2\pi} a)^{-1} \cdot \exp\big[-\theta^2/2a^2\big], \quad \mathsf{x} = \mathsf{rec}, \mathsf{proj}, \tag{S2}$$

where $a$ controls the width of the Gaussian curve. Given this Gaussian connection, it can be verified that neural responses of the attractor states are expressed as follow:

$$\bar{u}^E(\theta, t) = \mathsf{A}_u \exp[-(\theta - z(t))^2/4a^2], \quad \bar{r}^E(\theta, t) = \mathsf{A}_r \exp[-(\theta - z(t))^2/2a^2], \tag{S3}$$

where $\bar{u}(\theta, t)$ and $\bar{r}(\theta, t)$ are both Gaussian curves centered at the encoded stimulus $z(t)$. Combining above equations, we can derive the following expressions:

$$\mathsf{A}_r = \frac{g(t) \cdot \mathsf{A}_u^2}{1 + w^{IE} \rho \sqrt{2\pi} a \mathsf{A}_u^2}, \tag{S4a}$$

$$r^I(t) = \rho \cdot \sqrt{2\pi} a \cdot \mathsf{A}_u^2. \tag{S4b}$$

##### S1.2 Up and down states by gain modulation

A significant feature of CAN is that its dynamics is dominated by a few motion modes, among which the basis function $\phi_0(\theta|z) = \exp[-(\theta - z)^2/4a^2]$ corresponds to the amplitude of the neural activity bump. By projecting Eq. (S1a) onto $\phi_0$, we derive the dynamics of $\mathsf{A}_u$:

$$\tau_u \dot{\mathsf{A}}_u(t) = -\mathsf{A}_u(t) + \frac{\rho w_{rec}^{EE}}{\sqrt{2}} \cdot \frac{g(t) \cdot \mathsf{A}_u(t)^2}{1 + k\rho\sqrt{2\pi} a \mathsf{A}_u(t)^2} = -\mathsf{A}_u(t) + g(t) \cdot f(\mathsf{A}_u(t)). \tag{S5}$$

As illustrated in Fig. 2E, $f(\cdot)$ is a S-shaped activation function. When the gain factor $g$ is large enough, a nonzero stable point will come into existence. A persistent neural activity can be hold on this stable point, which corresponds to the up state of the network. As $g$ decreases, the stable point will move leftward until there is no intersection between $\mathsf{A}_u$ and $f(\mathsf{A}_u)$, then the network drops to the down state. By modifying gain factor $g$, the network can be altered between up and down states.

#### S2 Information routing between slots

As stated above, a CAN can be switched between up and down states by gain modulation, which alters the ability of storing and passing information of the network. In the up state, the network is easy to be activated so the input stimulus will be held and passed on. In contrast, if it is in the down state, the network will not respond to the input and the information flow will be blocked. Based on this property, we can use gain modulation to dynamically route the information flow between memory slots.

Specifically, we applied two temporary slots between the two memory slots, and a command neuron in charge of the swapping procedure. The temporary slots are in down states by default, and can only be set into up state by gain modulation from the command neuron. Once gained, a temporary slot will be activated by the inputs from the memory slot and transmit the inputs to the other memory slot it projects to. Negative feedback will shut down the command neuron once the transmission is completed. Overall, the stimulus stored in the memory slots will be swapped mediated by the temporary slots under the gain modulation of command neuron. Slots are attenuated by mutual inhibition to ensure the input and output information will not cause conflicts.

##### S2.1 Command neuron

We use a separate neuron to transmit the swap command to the memory and temporary slots. Denote its synaptic input and firing rate as $u_c$ and $r_c$ respectively, the neuronal dynamics will write:

$$\tau_c \dot{u}_c(t) = -u_c(t) + w_c \cdot r_c(t) + I_{cue} - I_{CM}, \tag{S6a}$$

$$r_c(t) = \frac{g_c(t) \cdot [u_c(t)]_+^2}{1 + k_c [u_c(t)]_+^2}, \tag{S6b}$$

$$g_c(t) = g_c^0 + I_G \tag{S6c}$$

Similar to Eq. (S5), the firing rate of the command neuron will remain high once activated by the cue signal $I_{cue}$. $I_{CM}$ is the negative feedback that we will define in the following subsections, and $I_G$ describes the gain modulation.

The gain of temporary slots will be modulated by the output of command neuron:

$$g(t) = g_0 + w_{TC} \cdot r_c(t), \tag{S7}$$

where $\mathsf{W}_{TC}$ denotes the weight from command neurons to temporary slots.

##### S2.2 Shared inhibition pool

Intuitively, each time a memory slot exports the stimulus that it encodes to a temporary slot, its activity should be eliminated for the upcoming new inputs, and vice versa for temporary to memory slots, which implicates a competition between them. Therefore, we introduced an inhibition pool, which renders a mutual inhibition. Specifically, besides the corresponding slot, the interneuron also accept input from the adjacent two slots. For example, the interneuron of memory slot $M_1$ receives inputs from temporary slots $T_{12}$ and $T_{21}$, and temporary slot $T_{12}$ receives from $M_1$ and $M_2$, which can be captured by the following equations:

$$\tau_I \dot{r}_m^I(t) = -r_m^I(t) + \rho \Big[ \int [u_m^E(\theta', t)]_+^2 \mathsf{d}\theta' + w_{mn}^{IE} \sum_n \int r_n(\theta'', t) \mathsf{d}\theta'' \Big], \tag{S8}$$
$$m, n \in \{M_1, M_2, T_{12}, T_{21}\}, \quad m \neq n,$$

where $w_{mn}^I E$ is the inhibitory connection strength between slot $m$ and $n$.

##### S2.3 Feedback

Since cues are usually transiently presented, the command neurons need an attractor dynamics to sustain the swap command by themselves. Therefore, after the swap has been executed successfully, a negative feedback will be needed to actively turn off the command neuron. The signature of completion is the reactivation of memory slots, so we assume the feedback strength is determined by the activity of memory slots:

$$I_{CM} = w_{CM} \cdot \rho \sum_m \Big[ \int r_m(\theta', t) \mathsf{d}\theta' \Big], \quad m \in \{M_1, M_2\}, \tag{S9}$$

where $w_{CM}$ is a parameter denoting the feedback strength.

### S3 Action composition

Given the implementation of swap operation, we can decompose an arbitrary permutation into sequential execution of swaps. By arranging them in a tree-like structure, we can efficiently implement the whole permutation group by combining the swaps in different orders.

#### S3.1 Permutation group

A *permutation* is an operation that maps a set of elements into itself by reordering their positions. If the set contains n elements, then the permutation is of degree n. Given two permutations $s, t$ of a set $A$, we can define the product $st$ as consecutive execution of $t$ and $s$ on $A$. With this definition, all permutations of the set $A$ constitute a *permutation group*, denoted as $S_n$.

Typically, a permutation can be expressed in a two-line notation. Consider a permutation $s$ that maps set $A$ from arrangement $(a_1, a_2, \cdots, a_n)$ to $(b_1, b_2, \cdots, b_n)$, then its two-line notation can be written as:

$$s = \begin{pmatrix} a_1, & a_2, & \cdots, & a_n \\ b_1, & b_2, & \cdots, & b_n \end{pmatrix}. \tag{S10}$$

##### S3.1.1 Cycle

A cycle is a special permutation which permutes the set in a cyclic manner. That is to say, a cycle $c$ of $m$ elements can be represented as $c = \begin{pmatrix} a_1, & a_2, \cdots, & a_{m-1}, & a_m \\ a_2, & a_3, \cdots, & a_m, & a_1 \end{pmatrix}$. For conciseness, cycle $c$ can be simplified as:

$$c = \Big(a_1, a_2, \cdots, a_{m-1}, a_m\Big), \tag{S11}$$

which is called cycle notation.

Two cycles are said to be disjoint if they have no elements in common. We have following lemma:

**Lemma 1** (Permutation Decomposition 1)**.** *Every permutation in the permutation group $S_n$ can be written as a product of disjoint cycles.*

*Proof.* Let $\sigma \in S_n$ be a permutation. To construct the cycle decomposition, begin with any element $a \in \{a_1, a_2, \cdots, a_n\}$. Apply $\sigma$ repeatedly to generate the sequence

$$a,\ \sigma(a),\ \sigma^2(a),\ \ldots$$

Since $\sigma$ acts on a finite set, this sequence must eventually repeat. Let $k$ be the smallest positive integer such that $\sigma^k(a) = a$. Then the set $\{a, \sigma(a), \ldots, \sigma^{k-1}(a)\}$ forms a cycle:

$$(a\ \sigma(a)\ \ldots\ \sigma^{k-1}(a)).$$

Remove these elements from consideration and repeat the process with another element not in the cycle, until all elements are exhausted. The result is a product of disjoint cycles. □

##### S3.1.2 Transposition

A transposition is a cycle of two elements, which swaps their positions. Any cycle can be decomposed into transpositions. We have the following lemma:

**Lemma 2** (Cycle Decomposition)**.** *Every cycle of length $k \geq 2$ can be written as a product of $k-1$ transpositions. That is, the cycle $(a_1\, a_2\, \ldots\, a_k)$ can be expressed as*

$$(a_1\, a_2\, \ldots\, a_k) = (a_1\, a_k)(a_1\, a_{k-1}) \cdots (a_1\, a_3)(a_1\, a_2).$$

*Proof.* The proof is omitted as it is straightforward.

□

Lemma.2 proved that all cycles can be decomposed into transpositions, but these transpositions do not always act on adjacent elements. By the following lemma, we can convert every transposition into a product of adjacent ones:

**Lemma 3** (Adjacent transposition). *For elements $e_1, e_2, e_k$, the following identity holds:*

$$(e_1, e_k) = (e_2, e_k)(e_1, e_2)(e_2, e_k).$$

*Proof.* The lemma can be simply proved by written step-by-step with intermediate states:

$$(e_2, e_k)(e_1, e_2)(e_2, e_k) \quad = \quad \begin{pmatrix} e_1 & e_2 & e_k \\ e_1 & e_k & e_2 \\ e_1 & e_k & e_2 \\ e_2 & e_k & e_1 \\ e_2 & e_k & e_1 \\ e_k & e_2 & e_1 \end{pmatrix} = (e_1, e_k)$$

Thus, we have:

$$(e_1, e_k) = (e_2, e_k)(e_1, e_2)(e_2, e_k).$$

□

Combining Lemma.1,2,3, it will be easy to conclude that any arbitrary permutation can be written as a product of adjacent transpositions. We conclude above statement into a theorem:

**Theorem 1** (Permutation Decomposition 2). *Every permutation in the permutation group $S_n$ can be written as a product of adjacent transpositions of the form $(a_i\ a_{i+1})$, where $1 \leq i < n$.*

*Proof.* Omitted. □

##### S3.2 $S_3$ group

$S_3$ group consists of all permutations of 3 items, namely, the set $\{1, 2, 3\}$. It contains 6 elements: $(), (1, 2), (2, 3), (1, 3), (1, 2, 3), (1, 3, 2)$, where $()$ denotes the identity element. Tab.S1 presents the Cayley table of $S_3$ group, indicating the results of group multiplication. Each entry of the table is the product of the column and row elements.

Table S1: Cayley table of the $S_3$ group

| **Element** | $()$ | $\mathbf{(1,2)}$ | $\mathbf{(2,3)}$ | $\mathbf{(1,3)}$ | $\mathbf{(1,2,3)}$ | $\mathbf{(1,3,2)}$ |
|---|---|---|---|---|---|---|
| $()$ | $()$ | $(1,2)$ | $(2,3)$ | $(1,3)$ | $(1,2,3)$ | $(1,3,2)$ |
| $\mathbf{(1,2)}$ | $(1,2)$ | $()$ | $(1,2,3)$ | $(1,3,2)$ | $(2,3)$ | $(1,3)$ |
| $\mathbf{(2,3)}$ | $(2,3)$ | $(1,3,2)$ | $()$ | $(1,2,3)$ | $(1,3)$ | $(1,2)$ |
| $\mathbf{(1,3)}$ | $(1,3)$ | $(1,2,3)$ | $(1,3,2)$ | $()$ | $(1,2)$ | $(2,3)$ |
| $\mathbf{(1,2,3)}$ | $(1,2,3)$ | $(1,3)$ | $(1,2)$ | $(2,3)$ | $(1,3,2)$ | $()$ |
| $\mathbf{(1,3,2)}$ | $(1,3,2)$ | $(2,3)$ | $(1,3)$ | $(1,2)$ | $()$ | $(1,2,3)$ |

##### S3.3 Tree-like control flow

By examining Tab.S1, we can naturally divide the $S_3$ group into 3 collections: $\{()\}, \{(1, 2), (2, 3)\}$ and $\{(1, 2, 3), (1, 3, 2), (1, 3)\}$, by whether they are identity element, adjacent transpositions and others. Elements in the last collection can be obtained by group multiplication of elements in the second one. This indicates a tree-like structure in terms of implementation, as we showcased in Sec. 4 (Fig. 4A). Specifically, we only need to implement the three transpositions in the circuit, then the 3-permutations can be performed by sequential execution of transpositions. This will require a tree-like control flow.

**Tree structure of command neurons** Take $(1, 3, 2) = (2, 3)(1, 2)$ as an example. We denote the corresponding command neurons as $\text{Cmd}_{(1,2)}, \text{Cmd}_{(2,3)}$ and $\text{Cmd}_{(1,3,2)}$, and arrange them in a tree structure, where $\text{Cmd}_{(1,3,2)}$ is posited at the upper level, dictating $\text{Cmd}_{(1,2)}$ and $\text{Cmd}_{(2,3)}$ at the lower level (Fig. 4A). The firing of these command neurons should satisfy three requirements: **1.** Activation of $\text{Cmd}_{(1,3,2)}$ will trigger the firing of $\text{Cmd}_{(2,3)}$ and $\text{Cmd}_{(1,2)}$. **2.** $\text{Cmd}_{(2,3)}$) and $\text{Cmd}_{(1,2)}$ should be activated in order, indicating the sequential initiation of the underlying swaps. **3.** There should be only one of the lower level command neurons firing at a time, suggesting a mutual inhibition.

Therefore, besides the inputs presented in Eq. (S6), $\text{Cmd}_{(2,3)}$ and $\text{Cmd}_{(1,2)}$ also receive an excitatory command inputs $I_{\mathsf{cmd}}$ from $\text{Cmd}_{(1,3,2)}$, a mutual inhibition input $I_{mut}$ from each other and a top down gain modulation input:

$$\begin{aligned} I_{\mathsf{cmd}} &= w_{\mathsf{cmd}} \cdot r_c^{(1,3,2)}, \\ I_{mut} &= w_{mut} \cdot r_c^l, \quad l = (1,2),(2,3), \\ I_G &= w_G^m \cdot r_c^{(1,3,2)}, \quad m = 1,2, \end{aligned}$$

where $w_{\mathsf{cmd}}$ denotes the input strength to $\text{Cmd}_{(2,3)}$ and $\text{Cmd}_{(1,2)}$. For the two neurons to be activated consecutively, we introduced a gradient on the gain input, in this case, $w_G^1 > w_G^2$.

The same as in swap, firing of lower level command neuron will be terminated by negative feedback after that the swap has been executed. Then command neurons with weaker input will be released from mutual inhibition and triggered to perform the following actions.

**Inhibition of return** Besides composition of two operations, there could be more of them in a broader scenario, for example, composition of operation $A, B, C$ into $ABC$, which requires a sequence of actions $C \to B \to A$. This setting will cause a nuanced problem, that is, reversal of the operation sequence. When the command neuron $\mathsf{Cmd}_B$ has been terminated, $\mathsf{Cmd}_C$ will become the one that receives the strongest input and most likely to fire, instead of the right one $\mathsf{Cmd}_A$. Therefore, we introduced a conjugate neuron for every command neuron to trace their firing history in the operation sequence (Fig. 4A). Their dynamics can be expressed as:

$$\tau_{\mathsf{cg}} \dot{u}_{\mathsf{cg}}^m(t) = -u_{\mathsf{cg}}^m(t) + w_{\mathsf{cg}} \cdot r_{\mathsf{cg}}^m(t) + I_{\mathsf{cg}}^E, \tag{S12a}$$

$$r_{\mathsf{cg}}^m(t) = \frac{g_{\mathsf{cg}}(t) \cdot [u_{\mathsf{cg}}^m(t)]_+^2}{1 + k_{\mathsf{cg}} \cdot [u_{\mathsf{cg}}^m(t)]_+^2}, \tag{S12b}$$

$$g_{\mathsf{cg}}(t) = g_{\mathsf{cg}}^0 + w_G^m \cdot r_c^{ABC}(t), \tag{S12c}$$

$$I_{\mathsf{cg}}^E = w_{\mathsf{cg}}^E \cdot r_c^m(t), \tag{S12d}$$

$$m = A, B, C. \tag{S12e}$$

Their dynamics is similar to that of command neurons but a subtle difference. When only one single operation is executed, it is not necessary to trace the history. Hence, $g_{cg}^0$ is set to not be able to hold a persistent activity by itself. Furthermore, conjugate neurons $\mathsf{Cg}$ for $A/B/C$ are gain modulated by command neuron $\mathsf{Cmd}_{ABC}$. Once gained, they will be triggered by the firing of its corresponding command neuron $\mathsf{Cmd}_{A/B/C}$, hold the history until the end of the whole operation sequence, and inhibit its $\mathsf{Cmd}$ from activated again. Following the completion of the operations, all conjugate neurons of all executed operations should be activated, and their activities together will be just the amount needed to terminate $\mathsf{Cmd}_{ABC}$. After the termination of $\mathsf{Cmd}_{ABC}$, the history traced by Cgs will also be cleared due to lack of gain input from it.

Hence, the overall dynamics of the $\mathsf{Cmd}$ neurons can be unfolded as:

$$\begin{aligned} \tau_c \dot{u}_c^m(t) =& - u_c^m(t) + w_c \cdot r_c(t) + w_{\mathsf{cmd}} \cdot r_c^l(t) - w_{mut} \cdot (\sum_m r_c^m(t)) \\ & - w_{\mathsf{cg}}^{fb} \cdot (\sum_n r_{\mathsf{cg}}^n(t)) - w_{\mathsf{cg}}^I \cdot r_{\mathsf{cg}}^m(t) + I_{cue} + I_{CM}, \end{aligned} \tag{S13a}$$

$$r_c^m(t) = \frac{g_c^m(t) \cdot [u_c^m(t)]_+^2}{1 + k_c^m [u_c^m(t)]_+^2}, \tag{S13b}$$

$$g_c^m(t) = g_c^0 + w_G^m \cdot r_c^l(t). \tag{S13c}$$

where $\text{level}(l) > \text{level}(m), \text{level}(n) < \text{level}(m)$ and $m = A, B, C, ABC$. The terms on the right-hand side of Eq. (S13a) are self-decay of command neuron, recurrent input from itself, top-down command input, mutual inhibition between command neurons within the same level, negative feedback from the lower level conjugate neurons, inhibition of return from the conjugate neuron of its own, external cue inputs and feedback from memory slots (if the command neuron is located on the lowest level).

#### S4 Motor readout

We introduced a motor module to readout the stimulus stored in the memory slots to generate behavioral responses. The neural response of the motor module can be described as:

$$\tau_{mo}\dot{r}_{mo}(t) = -r_{mo}(t) + [\mathsf{W}_{out}\sum_{m} r^{m}(t) - \Theta], \tag{S14}$$
$$r_{mo} = [r_{mo}]_{+}, \quad m = M_1, M_2, M_3, \cdots ,$$

where $\Theta$ is the readout threshold. Before cued to respond, the memory slot activities will always remain below $\Theta$. The order of readout is controlled by a timing circuit, which forms a chain structure, by projecting gain input to memory slots from different segments on the chain. The autonomous dynamics of the timing chain will ensure that, once initiated by the 'go' cue, the neural activity will propagate along the chain, rendering the timing and order of certain slots. The timing chain is implemented as a CAN, but the recurrent connection is slightly skewed to introduce a motion bias:

$$\tau_{tm}\dot{u}_{tm}(\theta, t) = -u(\theta, t) + \rho\mathsf{W}_{tm}(\theta) \star r_{tm}(\theta, t) + I_{go}, \tag{S15a}$$

$$r_{tm}(\theta, t) = \frac{[u_{tm}(\theta, t)]_{+}^{2}}{1 + k_{tm}\cdot\rho\int[u_{tm}(\theta', t)]_{+}^{2}\mathsf{d}\theta'}, \tag{S15b}$$

$$\mathsf{W}_{tm}(\theta) = w_{tm}^{0}\exp[-\theta^2/2a^2] - w_{tm}^{skew}\theta\exp[-\theta^2/2a^2], \tag{S15c}$$

where the recurrent connection $\mathsf{W}_{tm}(\theta)$ is the sum of a symmetric and an asymmetric term. $I_{go}$ is an input targeting the beginning of the chain when 'go' cue is presented:

$$I_{go} = \delta(t - t_{go}) \cdot I_{go}^{0}\exp[-(\theta - 0)^2/4a^2]. \tag{S16}$$

$t_{go}$ denotes the time when 'go' cue is presented. An activity bump will be triggered at the beginning and travel along the chain. Memory slots receiving gain input from the chain, which can be expressed as:

$$g^{m}(t) = g_{0}^{m} + \int \mathsf{W}_{G}^{m}(\theta - \theta^{m}) \cdot r_{tm}(\theta, t)\mathsf{d}\theta, \quad m = M1, M2, M3\cdots \tag{S17}$$

where $\mathsf{W}_{G}^{m}(\theta - \theta^{m}) = w_{G}^{0}\exp[-(\theta - \theta^{m})^2/2a^2]$ is also a Gaussian function and $\theta^{m}$ denotes the corresponding point of slot $m$ on the chain. Therefore, when the timing chain travel through $\theta^{m}$, slot $m$ will be gained and exceed the threshold $\Theta$, and load the memory into the motor module.

#### S5 Simulation details

At the beginning of the simulation, stimuli are fed into the memory slots in the form of $I(\theta - z) = I_0\exp[-(\theta - z)^2/2a^2]$, where $z$ represents the position of the stimulus. To measure whether a stimulus is encoded by a memory slot, we calculate the similarity between the stimulus and neural response of the slot (3B):

$$s(t) = \langle r(\theta, t), I(\theta - z)\rangle / I_0, \tag{S18}$$

which is normalized over stimulus but preserve the effect of neural firing rate, indicating how close between their center and how strongly the neural response is encoding the stimulus (firing rate). $\langle \mathbf{x} \cdot \mathbf{y} \rangle$ denotes inner product of vector **x** and **y**.

The encoded stimuli $z(t)$ can also be linearly read out from the slots' response, $r(\theta, t)$ by using the population vector ([1]), i.e.,

$$z(t) = \text{Angle}\Big[\sum_{j} r(\theta_j, t)e^{i\theta_j}\Big], \tag{S19}$$

where $i = \sqrt{-1}$ is the pure imaginary number.

The setting of network parameters can be found in Table S2.

The code was written with *BrainPy* ([2]) and was simulated on MacBookPro laptop which has a M1 pro CPU and 32GB RAM.

#### S6 Supplementary figures

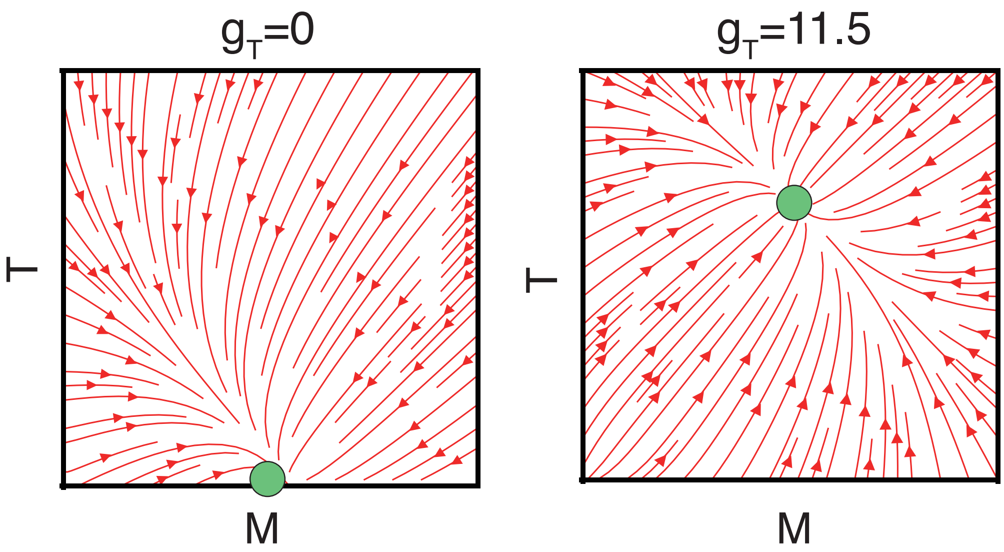

Figure S1: Vector field of mean firing rate of memory and temporary slots. Left: when gain of temporary slot $g_T = 0$, memory slot will win over temporary slot. Right: when $g_T$ is large, temporary slot will win and $r_T > r_M$.

#### References

[1] Apostolos P Georgopoulos, Andrew B Schwartz, and Ronald E Kettner. Neuronal population coding of movement direction. *Science*, 233(4771):1416–1419, 1986.

[2] Chaoming Wang, Tianqiu Zhang, Xiaoyu Chen, Sichao He, Shangyang Li, and Si Wu. Brainpy, a flexible, integrative, efficient, and extensible framework for general-purpose brain dynamics programming. *Elife*, 12:e86365, 2023.

Table S2: Typical parameters of the network model

| Symbol | Description | Values |
|---|---|---|
| $N^E$ | Number of neurons in one slot | 128 |
| $\rho$ | Neuronal density in the stimulus space | $N/2\pi$ |
| $a$ | Tuning width (rad) | 0.4 |
| $g_0$ | Baseline gain factor of slots | 0 |
| $g_c^0$ | Baseline gain factor of command neurons | 10 |
| $g_{\mathsf{cg}}^0$ | Baseline gain factor of conjugate neurons | 2 |
| $k$ | Inhibition strength of slots | 10 |
| $k_c$ | Inhibition strength of command neurons | 10 |
| $k_{cg}$ | Inhibition strength of conjugate neurons | 2 |
| $k_{tm}$ | Inhibition strength of timimg chain | 1 |
| $\tau$ | Synaptic decaying time constant of slots | 10 |
| $\tau_I$ | Synaptic decaying time constant of inhibitory neurons | 1 |
| $\tau_c$ | Synaptic decaying time constant of command neurons | 100 |
| $\tau_{cg}$ | Synaptic decaying time constant of conjugate neurons | 100 |
| $\tau_{mo}$ | Synaptic decaying time constant of motor module | 100 |
| $\tau_{tm}$ | Synaptic decaying time constant of timing chain | 1 |
| $\mathsf{d}t$ | Time step in numerical simulation | 0.1 |
| $w_{rec}^{EE}$ | Peak recurrent weight of slots | 1 |
| $w_{proj}^{EE}$ | Peak projection weight between slots | 0.5 |
| $w_{TC}$ | Gain modulation strength from command to slots | 12 |
| $w_{CM}$ | Feedback strength from memory slots to command neuron | 0.5 |
| $w^{IE}$ | Weight from E to I neurons in slots | 9.6 |
| $w_c$ | Recurrent weight of command neurons | 1. |
| $w_{cg}$ | Recurrent weight of conjugate neurons | 1. |
| $w_{mut}$ | Mutual inhibition strength between command neurons | 12 |
| $w_{\mathsf{cmd}}$ | Weight of E input from up to low level in control tree | 1.5 |
| $w_{cg}^E$ | Weight of E input from command neuron to conjugate neuron | 0.2 |
| $w_{cg}^I$ | Weight of I input from conjugate neuron to command neuron | 0.8 |
| $w_{cg}^{fb}$ | Weight of I feedback from low level conjugate neuron to high level command neuron | 0.05 |
| $w_G^1, w_G^2$ | Gain modulation strength from upper to lower control circuits | 3.3, 3.0 |
| $w_{tm}^0, w_{tm}^{skew}$ | Symmetric and asymmetric recurrent connection strength of timing chain | 1.5, 0.075 |
| $w_G^0$ | Weight of gain input from timing chain to slots | 100 |
| $\mathsf{W}_{out}$ | Readout weight | 1 |
| $\Theta$ | Readout threshold | 0.5 |
